# GPT-3 reveals selective insensitivity to global *vs.* local linguistic context in speech produced by treatment-naïve patients with positive thought disorder

**DOI:** 10.1101/2024.07.08.602512

**Authors:** Victoria Sharpe, Michael Mackinley, Samer Nour Eddine, Lin Wang, Lena Palaniyappan, Gina R. Kuperberg

**Affiliations:** Department of Psychology, Tufts University; Lawson Health Research Institute; Massachusetts General Hospital, Harvard Medical School; Robarts Research Institute, Western University, Ontario; Douglas Mental Health University Institute, McGill University

**Keywords:** lexical predictability, schizophrenia, large language models, computational psychiatry, natural language processing, context, language production

## Abstract

**Background:** Early psychopathologists proposed that certain features of positive thought disorder, the disorganized language output produced by some people with schizophrenia, suggest an insensitivity to global, relative to local, discourse context. This idea has received support from carefully controlled psycholinguistic studies in language comprehension. In language production, researchers have so far remained reliant on subjective qualitative rating scales to assess and understand speech disorganization. Now, however, recent advances in large language models mean that it is possible to quantify sensitivity to global and local context objectively by probing lexical probability (the predictability of a word given its preceding context) during natural language production.

**Methods:** For each word in speech produced by 60 first-episode psychosis patients and 35 healthy, demographically-matched controls, we extracted lexical probabilities from GPT-3 based on contexts that ranged from very local— a single preceding word: P(Wn | Wn-1)—to global— up to 50 preceding words: P(Wn|Wn-50, Wn-49, …, Wn-1).

**Results:** We show, for the first time, that disorganized speech is characterized by disproportionate insensitivity to global, versus local, linguistic context. Critically, this global-versus-local insensitivity selectively predicted clinical ratings of positive thought disorder, above and beyond overall symptom severity. There was no evidence of a relationship with negative thought disorder (impoverishment).

**Conclusions:** We provide an automated, interpretable measure that can potentially be used to quantify speech disorganization in schizophrenia. Our findings directly link the clinical phenomenology of thought disorder to neurocognitive constructs that are grounded in psycholinguistic theory and neurobiology.

## Introduction

Since the days of Kraepelin and Bleuler, psychopathologists have struggled to describe and understand the disorganized, incoherent language output produced by some people with schizophrenia — positive thought disorder (1,2). Positive thought disorder affects up to 50% of patients with schizophrenia (3) and is linked to significant impairments in social functioning (4,5) and overall quality of life (3,6). It is therefore of critical clinical importance to understand how to objectively quantify positive thought disorder, and to link this characterization to its underlying mechanisms.

One clue comes from careful descriptions of positive thought disorder; phenomena like tangentiality and derailment may indicate a relative insensitivity to global discourse context. Importantly, this insensitivity seems to be *specific* to global information, with sensitivity to local context being largely intact. In line with this idea, studies across multiple domains have documented reduced sensitivity to global, relative to local, context, in schizophrenia (7–10). Indeed, in language *comprehension*, several tightly controlled psycholinguistic studies, using both behavioral and neural measures, suggest that patients are less able than healthy controls to use global linguistic context (long sentences or discourse) to facilitate the processing of incoming words (11–15), whereas the automatic use of local linguistic context (e.g. directly related semantic primes, short sentence frames) is generally spared (13,16–20). In particular, Swaab and colleagues (13) showed that whereas healthy controls produced a larger N400 (an event-related potential that indexes the probability of words based on their context) in response to discourse-incongruent *versus* discourse-congruent words, people with schizophrenia only showed a congruency effect when the target word was predictable based on its local context.

Until now, however, it has not been possible to objectively quantify the use of global versus local context in natural speech *production* in schizophrenia. In the 1960s and 70s, some researchers attempted to assess patients’ general sensitivity to context using fill-in-the-blank “Cloze” completion tasks (e.g., (21–23)), but this procedure was extremely time-consuming and thus impossible to carry out on a large scale. It was also ill-suited for understanding patients’ *relative* sensitivity to global versus local context. Therefore, over 110 years after Bleuler’s seminal descriptions of disorganized speech in schizophrenia (1), clinicians and researchers remain reliant on subjective qualitative rating scales to assess and understand speech disorganization (positive thought disorder) in schizophrenia (24–30).

Fortunately, due to recent advances in Large Language Models (LLMs), it is now possible to obtain an automated, precise measure that objectively quantifies the relationship between each word and its full (local and global) preceding context — its lexical probability; that is the probability of observing that word, given the full set of words that precedes it: P(w_n_ | w_1_, w_2_, …, w_n-1_), where w_1_, w_2_, …, w_n_ is a sequence of words. *Lexical probability* is the most robust predictor of behavior (31,32) and neural activity (33–38) during language comprehension (for review, see (39)), and it is tightly linked to the coherence of language output (40,41 16410, 42–44). Indeed, although not typically designed with biological plausibility in mind, many large language models, like OpenAI’s GPT series, are explicitly trained to predict upcoming words based on their preceding context, and in doing so, they learn to produce coherent language that is remarkably human-like (45,46). Moreover, with certain manipulations to GPT’s use of context, it produces disorganized, incoherent speech that is very similar to that produced by people with positive thought disorder (47).

Over the past 15 years, various Natural Language Processing (NLP) measures have been used to achieve high classification accuracy when discriminating between schizophrenia patients and healthy controls--e.g., (48–54); see also (55) for an excellent overview--and predicting psychosis onset in clinical high-risk populations--e.g., (56,52,57 16061)).

While a few of these measures have been shown to correlate with clinical ratings of atypical speech--e.g., (48,58,59)--the degree to which these measures are linked *selectively* to *positive* thought disorder (i.e. language disorganization), as opposed to negative thought disorder (reduced overall production or impoverishment), has been largely unclear (though see (58) for work in clinical high-risk individuals). It is also unclear whether these measures account for variance in thought disorder above and beyond overall symptom severity, or medication use. Therefore, what is badly needed is a selective measure of positive thought disorder that is explicitly grounded in neurocognitive theory and that can not only help us *describe* positive thought disorder, but also *understand* its underlying cognitive and computational mechanisms.

In the present study, we build on this large NLP literature to develop such a measure. We used GPT-3, a state-of-the-art LLM, to estimate the lexical probability of every word in speech produced by untreated first-episode psychosis patients and demographically-matched healthy controls. Critically, we manipulated the length of the context window that the language model had access to, allowing us to obtain graded estimates of predictability for each word based on very local context—a single preceding word: P(W_n_ | W_n-1_)—to very global contexts—up to 50 preceding words: P(W_n_|W_n-50_, W_n-49_, …, W_n-1_). To anticipate our findings, we show for the first time that a disproportionate insensitivity to global versus local context specifically and selectively predicts clinical ratings of positive thought disorder. Thus, we provide an automated, theoretically-grounded measure that links clinical characterizations of thought disorder with a mechanistic understanding of language production.

## Methods

### Participants

One-hundred-and-six English-speaking participants (36 healthy controls: HCs; 70 first episode psychosis patients: FEPs) were included from an ongoing study (ClinicalTrials.gov Identifier: NCT02882204). All participants were between 16 and 45 years. Exclusion criteria were: a history of drug or alcohol dependence over the past year, a history of major head injury (with a period of unconsciousness or seizures), intellectual disability, uncontrolled medical conditions, >2 weeks of lifetime antipsychotic exposure, or the inability to provide informed consent.

FEPs were recruited April 2017 – September 2019 by screening all consecutive referrals to the Prevention and Early Intervention for Psychosis Program at the London Health Sciences Centre in London, Ontario, Canada. Patients were approached within two weeks of referral, ensuring that all were in the acute, untreated^1^ phase of psychosis. A later six-month consensus diagnosis from two research psychiatrists and the primary treatment provider, based on (61), and the Structured Clinical Interview for DSM-5 (62) indicated that 65 of these participants met criteria for a schizophrenia spectrum disorder and 5 for affective psychosis. The Research Ethics Board at Western University approved all study procedures and all participants provided informed consent.

Ten patients and one control participant were subsequently excluded from analysis because of uncertain Parental Socioeconomic Status (PSES (63)) information, which served as an important control variable (see below)^2^. Summary data of the 35 HCs and 60 FEPs included in the reported analyses are shown in Table 1.

**Table 1.**
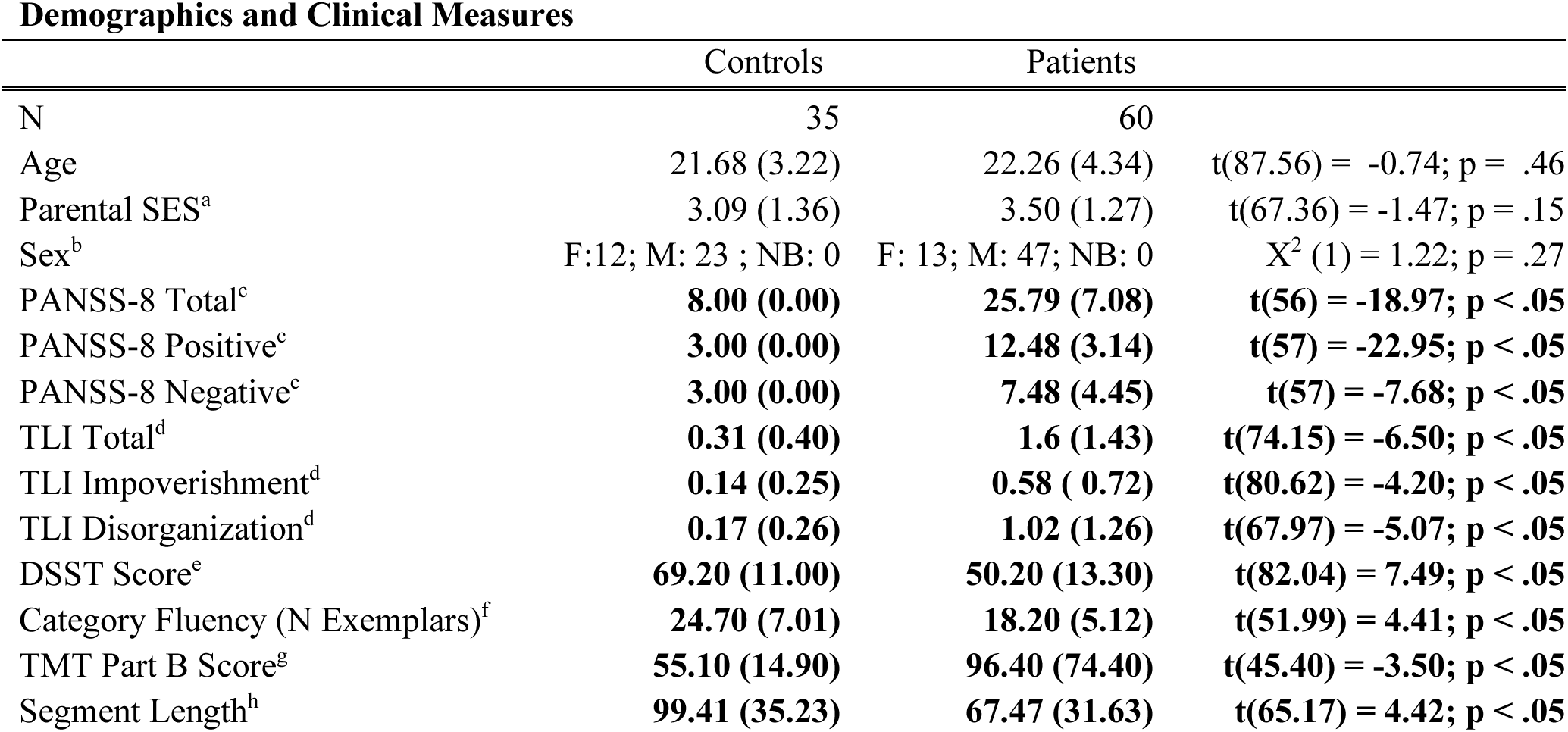
By-group means with standard deviations in parentheses; significant differences between groups shown in bold. ^a^(63); ^b^F = Female; M = Male; NB = Non-Binary/Intersex; ^c^(64,65); ^d^(30); ^e^Digit-Symbol Substitution Test (66), mean of Written and Oral scores; ^f^Category/Semantic Fluency: number of animal exemplars produced; ^g^Trail-Making Test, Part B (68); ^h^Mean length of contiguous speech segment (in words)

### Clinical and neuropsychological assessment

Patients’ symptoms were assessed using the 8-Item Positive and Negative Syndrome Scale (PANSS-8 (64); condensed from the PANSS (65) by one of two research psychiatrists, on the same week as speech acquisition (intraclass correlation for total scores (ICC) between the 2 raters; 10 subjects = 0.91).

In all participants, general cognitive function was assessed using three cognitive tasks: (1) the Digit-Symbol Substitution Test (DSST (66)) as a measure of working memory and processing speed; (2) the Category Fluency Task (67) as a measure of semantic memory and executive function, and (3) Part B of the Trail-Making Test (TMT (68)) as a measure of non-verbal executive function. See Supplementary Materials for details.

### Speech data

All participants described three pictures from the Thematic Apperception Test (69), for one minute each (see Supplementary Materials for details). Their speech was recorded and transcribed, and used for both clinical assessment of thought disorder and extraction of GPT lexical probability measures.

#### Thought disorder ratings

Thought disorder ratings were completed by a single trained graduate-level research assistant under the supervision of a research psychiatrist, blinded to patient status by using numbered transcripts, using the Thought and Language Index (TLI (30)). For each participant, the rater computed a measure of positive thought disorder ( “Disorganization” score) and a measure of negative thought disorder (“Impoverishment” score); see Supplementary Materials for details.

#### GPT-derived measures of lexical probability

OpenAI’s GPT-3 is a LLM with billions of parameters that can provide accurate estimates of the conditional probability of any given word (45), given a sequence of *k* preceding words (the preceding context), i.e. P(w_n_ | w_n-k_, w_n-k+1_, …., w_n-1_). We extracted the lexical probability of each word in each participant’s speech, based on all available context, as well as based on different context conditions (see Results; Figure 1). These extracted values served as the dependent measure in a series of linear mixed effects regressions designed to test our a priori hypothesis. To extract these various estimates of lexical probability, we used the ‘davinci_002’ model via the OpenAI API (https://openai.com/product), after splitting each participant’s transcript into contiguous discourse “segments.” Within each segment, spellings were standardized and all punctuation except for apostrophes, hyphens, and sentence-final punctuation was removed.

**Figure 1.**
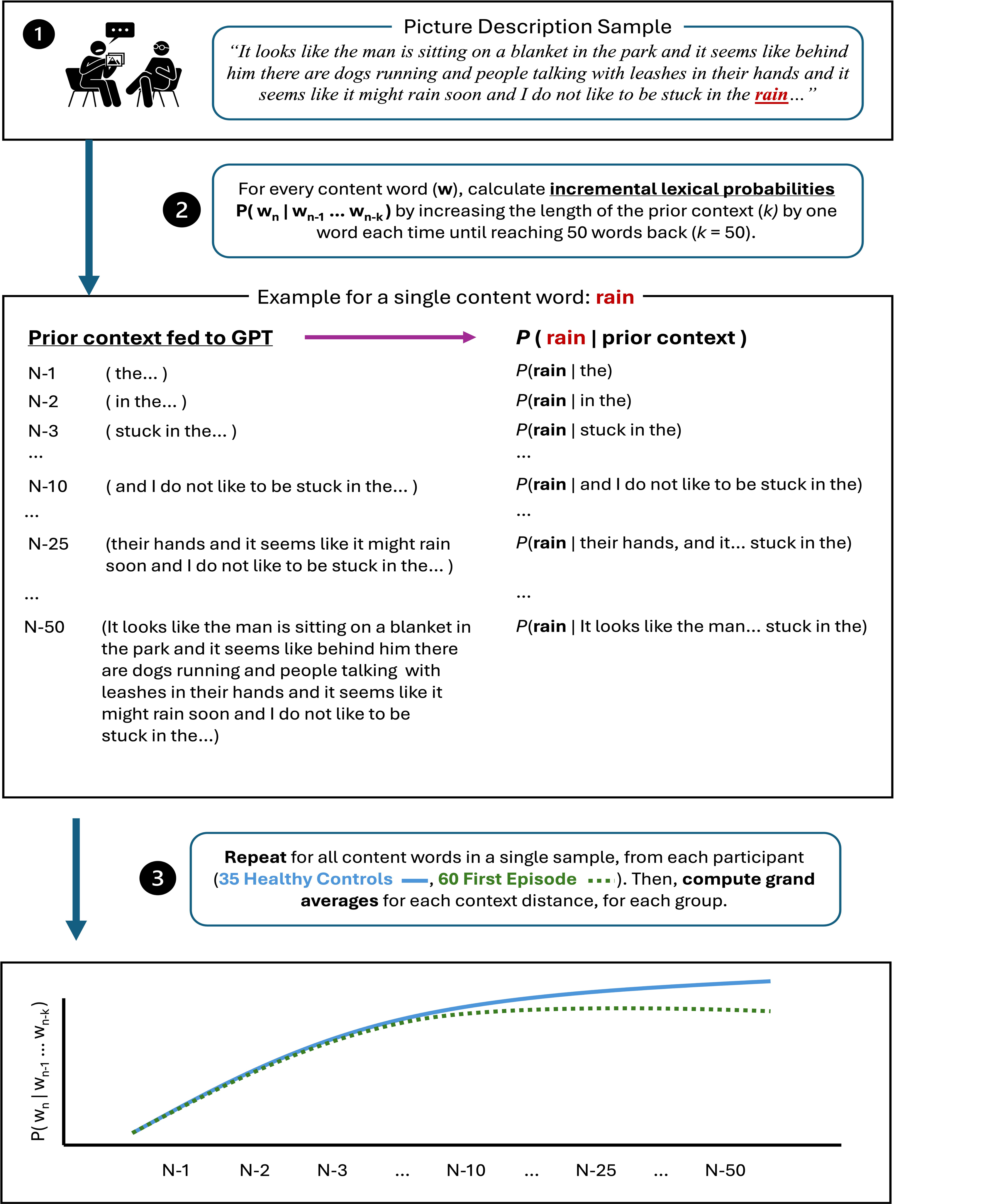
Process for extracting GPT lexical probabilities from participant speech samples given varying amounts of context. (*1*) Speech is transcribed, and spellings and punctuation are standardized. (*2*) For a given word *w*, we compute its lexical probability given 50 different context lengths, ranging from one word P(*w_n_* | *w_n-1_*) to 50 words P(*w_n_* | *w_n-50_*). (*3*) We repeat these computations for each word in each contiguous speech segment for each participant. These probabilities can then be used as single datapoints in mixed effects linear regression models, or averaged across context window sizes and groups, as in Figure 3.

### Statistical analysis

Prior to statistical analysis, we excluded any probability values for disfluencies (e.g., “um”, “uh”) and function words. This left a total of 19,421 word tokens (47.72% of the data; Patients: 11,696 tokens, 47.55%; Controls: 7725 tokens, 47.98%). We log-transformed all probability values to ensure that the assumptions of the general linear model were met and to allow us to probe proportional, rather than absolute, differences in lexical probability. We then excluded log probability values outside three standard deviations from the mean of each context condition, which resulted in the exclusion of 81.10 (SD = 15.36) datapoints (i.e. probability values) per condition (see above) on average.

Linear mixed effects regression provides a particularly advantageous analytic approach for this dataset, as it allowed us to test the effects of predictors of interest on Lexical Probability, while accounting for clustering in the data by incorporating by-subject and by-item (word token) random effects. It also allowed us to control for potential confounds by including “nuisance” item-level and participant-level covariates in each model: Segment Length (number of words in each segment), based on the possibility that participants distribute information differently across discourse segments of different lengths (42–44), and Parental SES (PSES; participant-level), which is associated with variance in various aspects of language use (see (70) for review). All continuous predictor variables were z-scored (see Supplementary Materials for details).

## Results

### 1. People with schizophrenia are less sensitive to context during language production

We began by asking whether FEPs were generally less sensitive to context than controls. We used GPT-3 to extract the lexical probability of each word in each discourse segment, based on the full set of words the participant had produced up until that point (the prior context)— P(Word | AllAvailableContext). Then, as a baseline, we extracted the lexical predictability of the same words, but replacing the prior context with randomly sampled words from an unrelated picture description (see Supplementary Materials)—P(Word | NoContext).

These lexical probabilities served as the outcome variable in a linear mixed effects model, with predictors of interest ContextType (AllAvailableContext vs. NoContext; within items), Group (FEPs vs. HCs; between participants), and the ContextType*Group interaction; see Table 2, Figure 2. As expected, across groups, participants produced words that were more predictable in the AllAvailableContext condition than in the NoContext condition (a main effect of ContextType). However, this effect was smaller in the patient group (an interaction between Context and Group), with follow-ups showing that, relative to HCs, FEPs produced less predictable words in the AllAvailableContext condition (Est.= -0.38, SE=0.10, p<.001), but not the NoContext condition (Est.=-0.07, SE =0.09, p=0.43).

**Figure 2.**
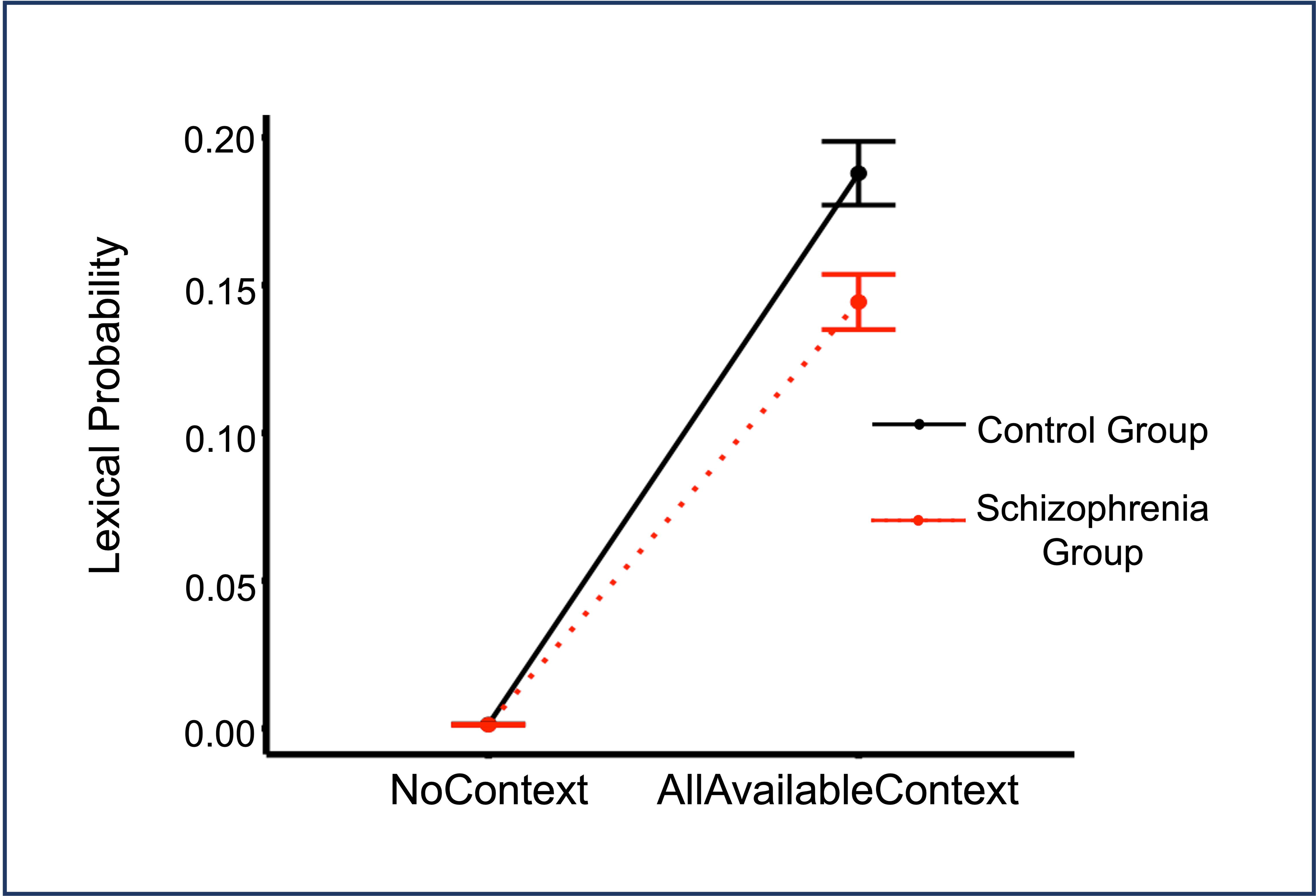
Mean lexical predictability, by context type (NoContext vs. AllAvailableContext) for controls vs. patients. Error bars represent standard error.

**Table 2.**
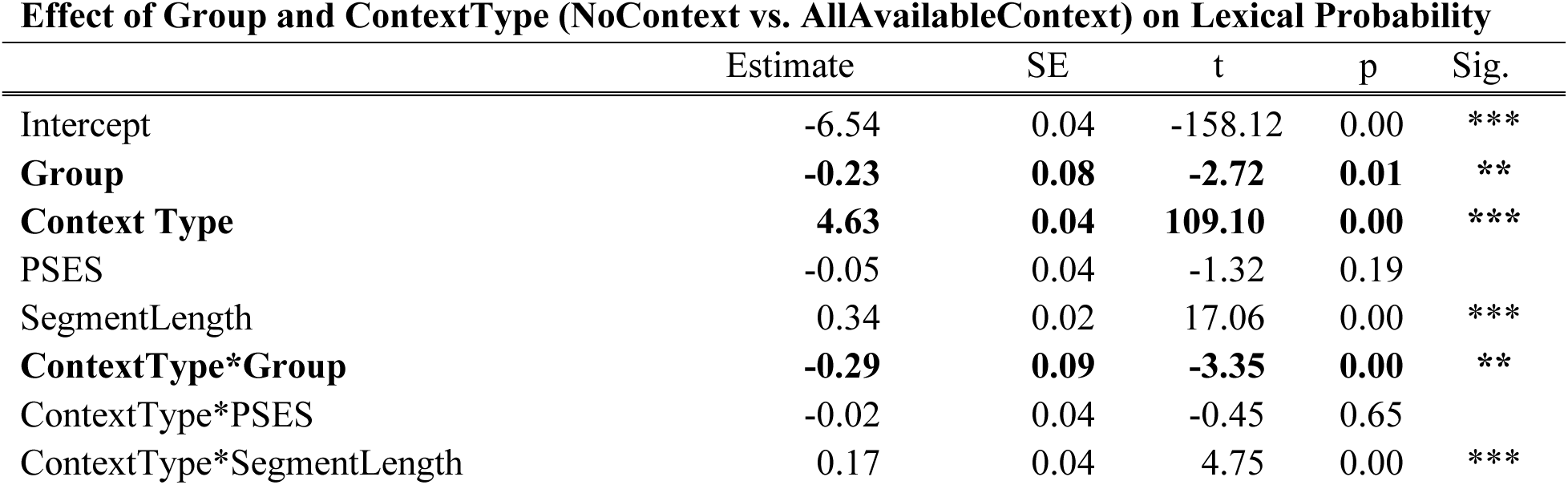
Predictors of interest are shown in bold.

### 2. People with schizophrenia exhibit a *selective* insensitivity to global (versus local) context

We next turned to the critical question of whether patients are selectively insensitive to global, relative to local, linguistic context.

For each word in each discourse segment, we extracted its lexical probability based on 50 different context window sizes (ranging from 1 preceding word to 50 preceding words; Figure 1), generating up to 50 unique datapoints for each word produced by each participant: 747,195 datapoints total. These values served as the outcome variable in an analysis where the predictors of interest were ContextWindowSize (continuous, within-items) and Group (FEPs vs. HCs; between-participants), and their interaction. ContextWindowSize was log-transformed to produce a more linear relationship with lexical probability (also log-transformed; see Methods), per the assumptions of the general linear model.

As expected, across both groups, larger ContextWindowSize predicted greater lexical probability (a main effect of ContextWindowSize). There was also a main effect of Group: at the mean ContextWindowSize (∼22.07 words of context), lexical probability was greater in HCs than in FEPs, consistent with the analysis above. Critically, however, as ContextWindowSize increased (i.e. as the length of the context available to the model increased), lexical probability increased *less* in patients, relative to controls (an interaction between ContextWindowSize and Group); Table 3, Figure 3.

**Figure 3.**
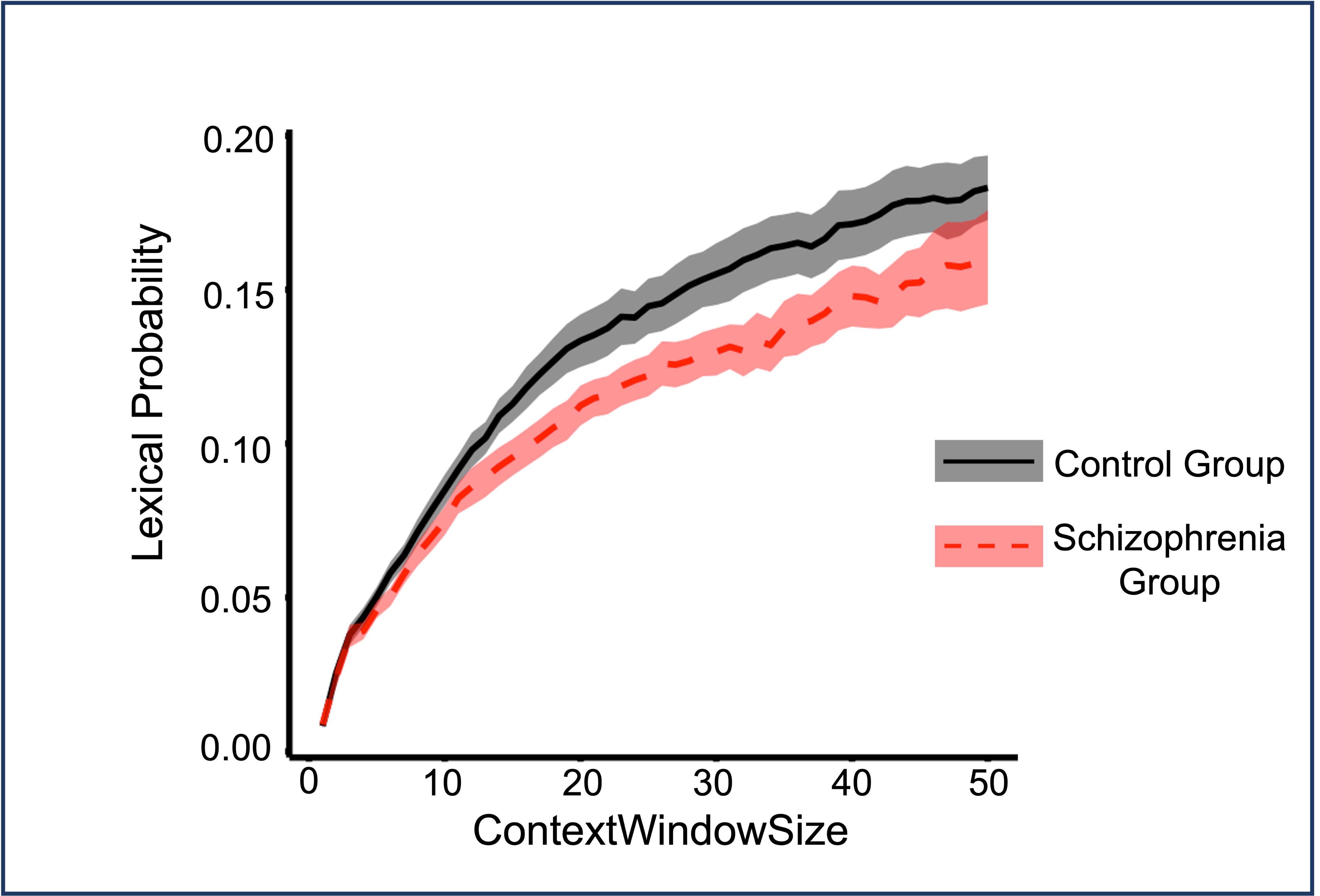
Mean lexical predictability, by ContextWindowSize (ranging from 1 to 50 words) for controls vs. patients. Shaded areas represent standard error.

**Table 3.**
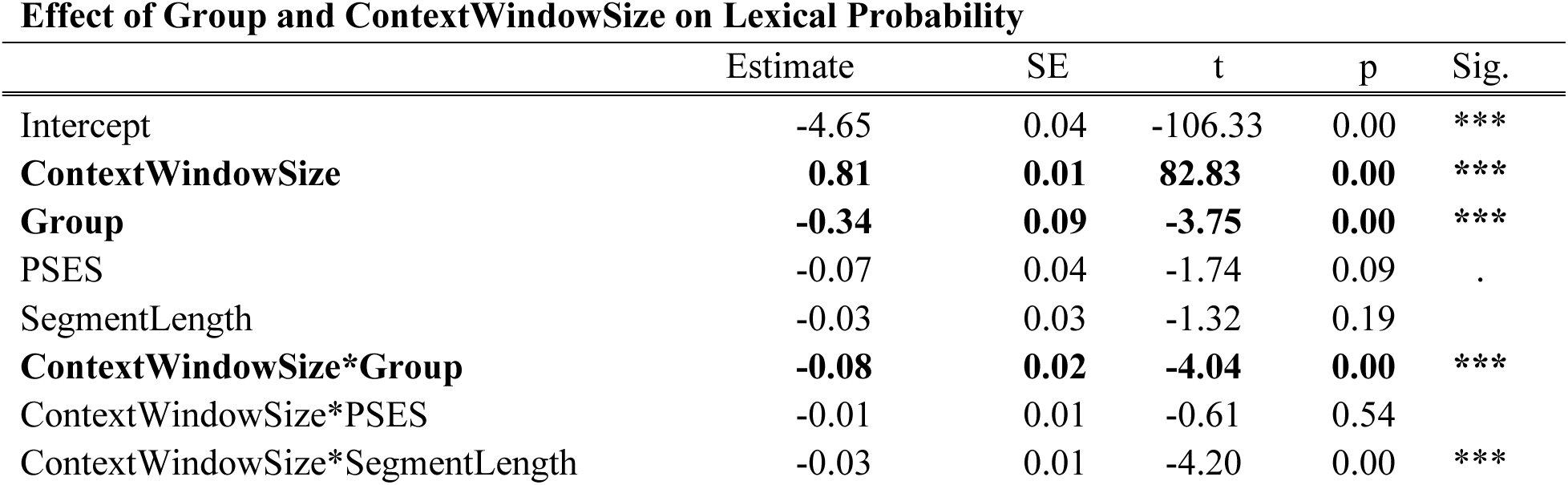
Predictors of interest are shown in bold.

Follow-ups at the local and global extremes of ContextWindowSize confirmed that for more global contexts (averaging across window sizes between 46 to 50 words), the group difference was significant (Est. = -0.25, SE = 0.09, p = 0.01), but for very local contexts (averaging across window sizes between 1-5 words), it was non-significant (Est. = -0.12, SE = 0.08, p = 0.13).

### 3. Relative insensitivity to global context is not driven by impairments in general cognitive function

To determine whether the interaction between Group and ContextWindowSize could be explained by differences in overall cognitive functioning, we averaged three scaled scores from each participant’s cognitive assessments (see Methods), and included this summary measure (CognitiveFunction), and its interaction with ContextWindowSize, as additional predictors in the above model.

The Group*ContextWindowSize interaction persisted, suggesting that selective insensitivity to global versus local context could not be explained by differences in CognitiveFunction between groups. Indeed, CognitiveFunction failed to predict lexical probability at all (no main effect of CognitiveFunction; no Group*CognitiveFunction interaction; Table 4)^3^.

**Table 4.**
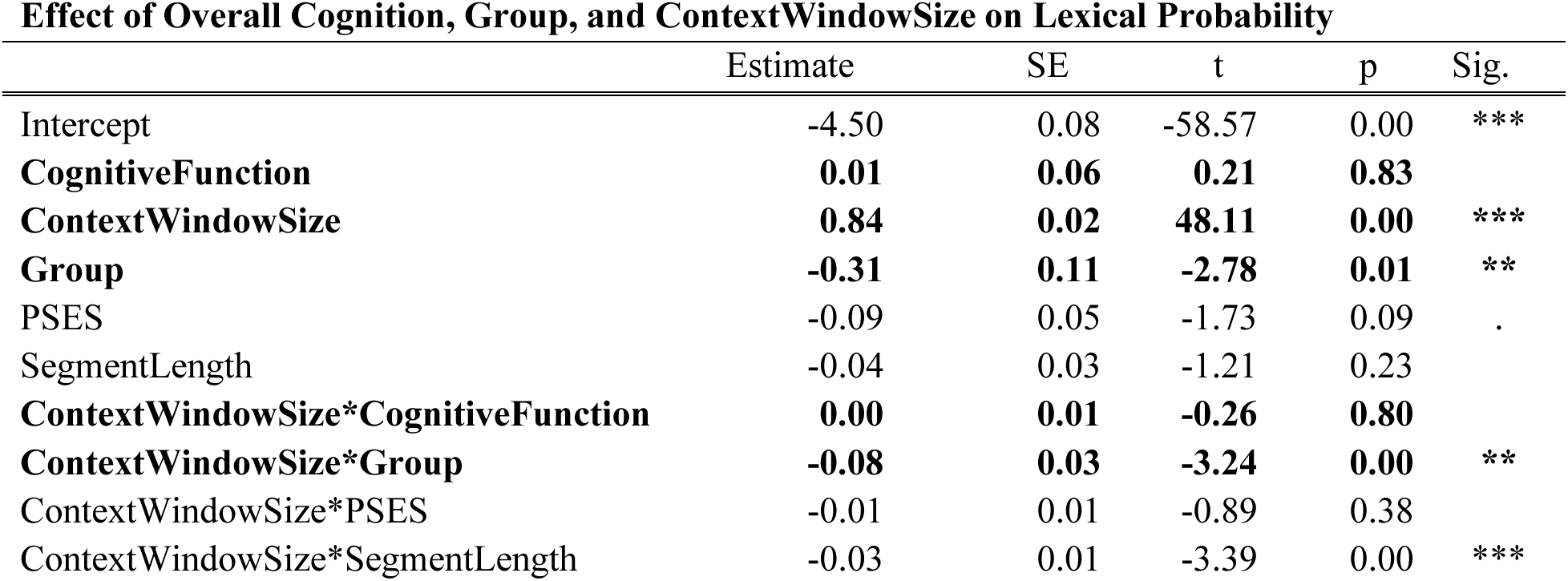
Predictors of interest are shown in bold.

### 4. Relative insensitivity to global context is selectively associated with positive thought disorder

We then asked whether, within the patient group, patients’ insensitivity to global context was linked to positive thought disorder. To address this question, we carried out an analysis, within the patient group only, with predictors of interest ContextWindowSize (continuous, within-items), Disorganization (TLI subscore; continuous, between-participants), and their interaction. This revealed a significant interaction between ContextWindowSize and Disorganization, such that Disorganization predicted the effect of ContextWindowSize on lexical probability. Specifically, greater Disorganization was associated with a smaller increase in lexical probability as ContextWindowSize increased; Table 5, Figure 4.

**Figure 4.**
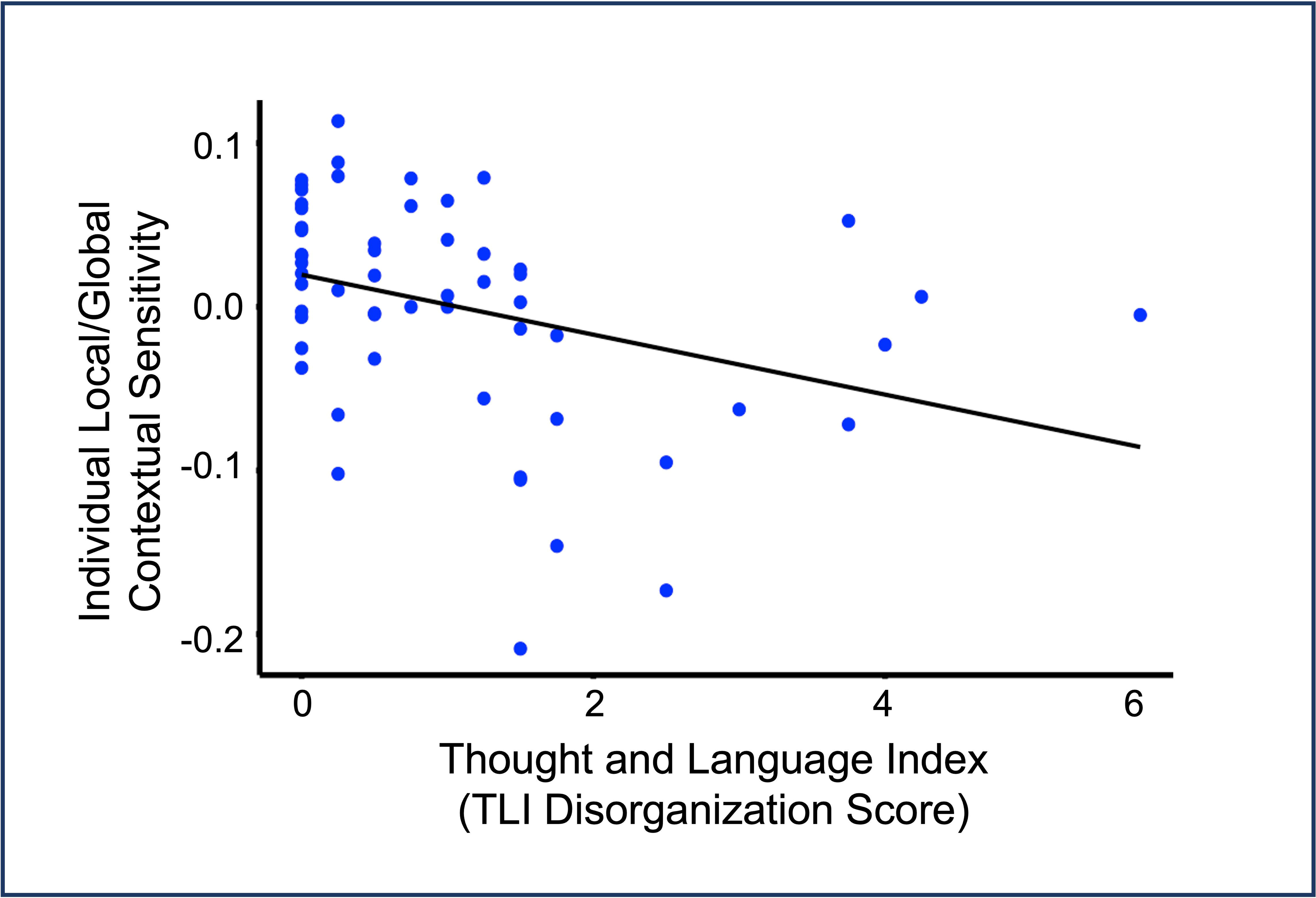
Relationship between speech disorganization, as measured by the TLI Disorganization sub-score (larger score indicates greater speech disorganization), and by-subject local/global bias (i.e. the by-subject slopes for the ContextWindowSize on lexical probability) within patients. Black line represents the regression line of best fit.

**Table 5.**
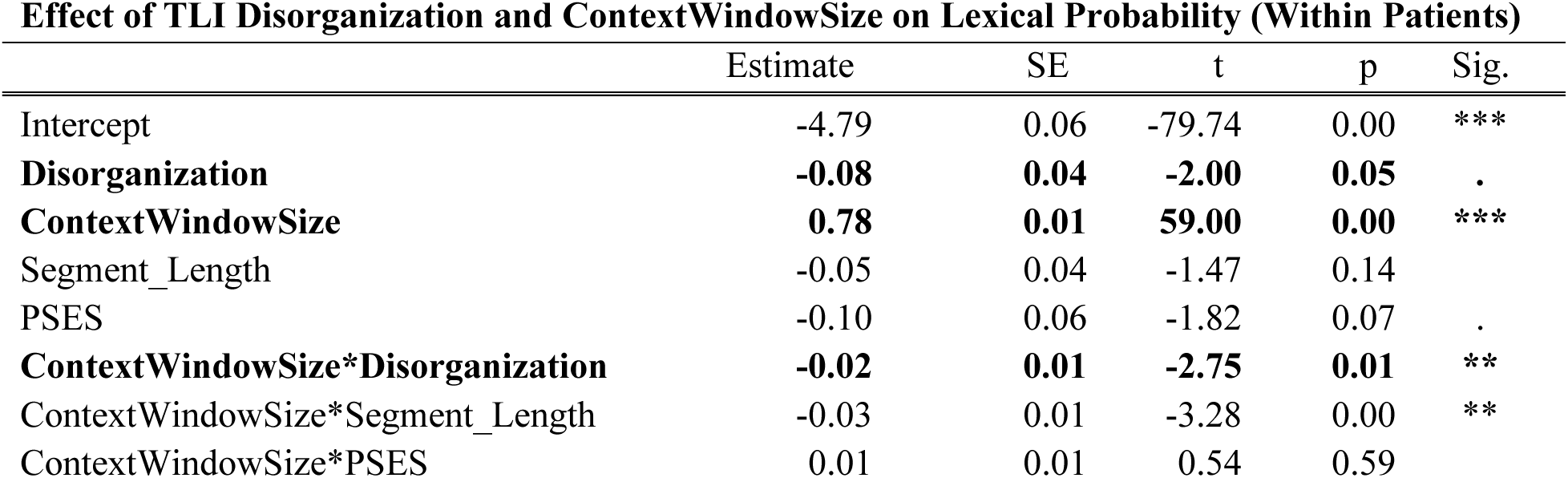
Predictors of interest are shown in bold.

Follow-ups at the local and global extremes of ContextWindowSize confirmed that for very local contexts (averaging across window sizes between 1-5 words), there was no effect of Disorganization (Est.=0.01, SE=0.04, p=.64), but for more global contexts (averaging across window sizes between 46 to 50 words), Disorganization significantly predicted lexical probability (Est.=-.11, SE=0.04, p=0.02).

To determine whether global-vs-local insensitivity was specifically linked to *positive* thought disorder, as opposed to negative thought disorder, we then ran an equivalent analysis in which we replaced Disorganization with the Impoverishment, as a measure of negative thought disorder. This revealed only a trending main effect of Impoverishment, and no Impoverishment*ContextWindowSize interaction; Table 6.

**Table 6.**
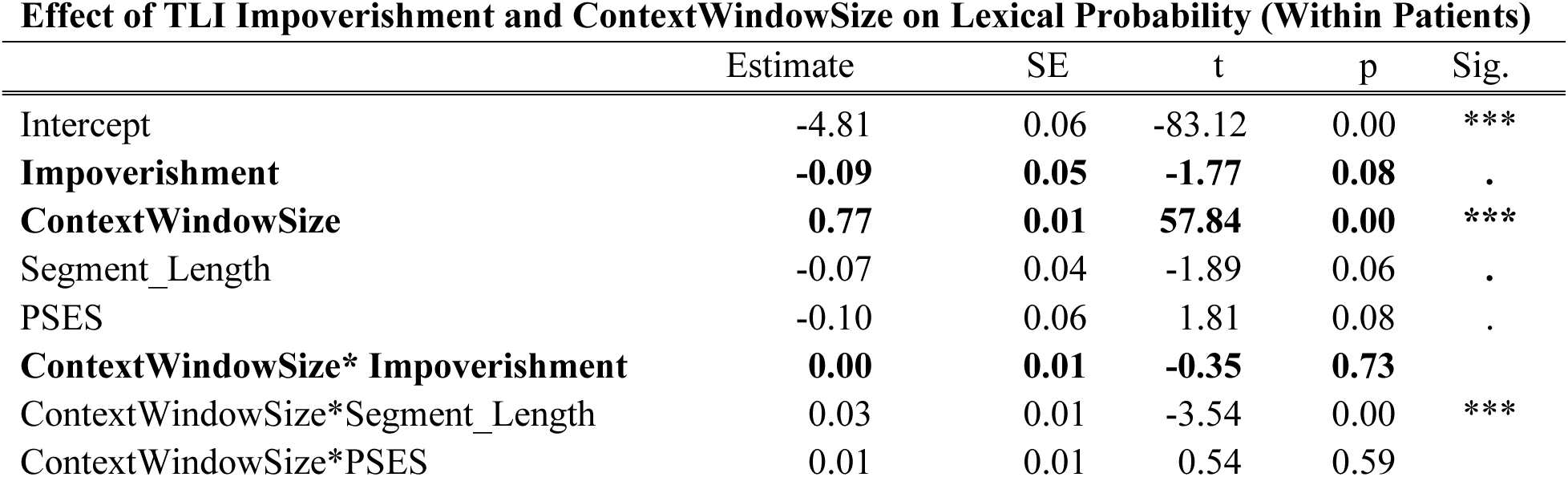
Predictors of interest are shown in bold.

Finally, to determine whether there was an effect of Disorganization over and above overall symptom severity, we ran an additional analysis which also included the PANSS-8 Total score and its interaction with ContextWindowSize (Table 7). The significant interaction between Disorganization and ContextWindowSize persisted. In contrast, neither the main effect of PANSS-8 nor the PANSS-8*ContextWindowSize interaction was significant.

**Table 7.**
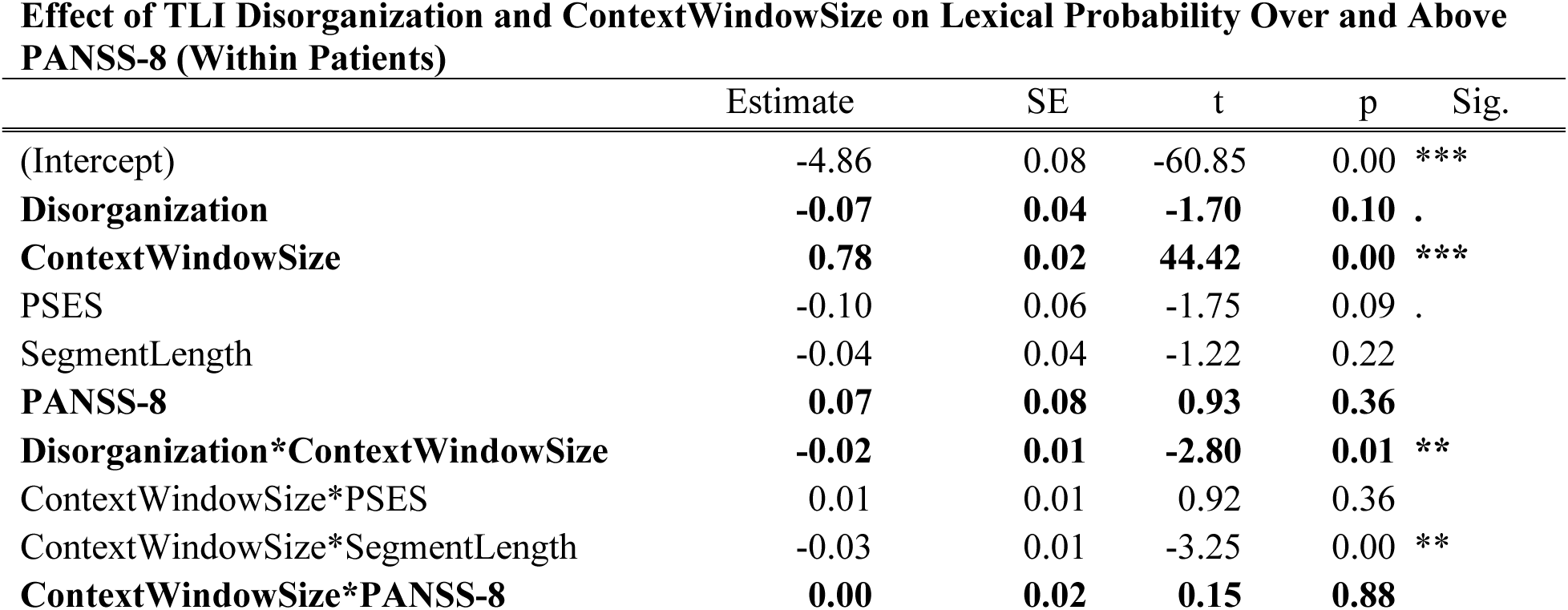
Predictors of interest are shown in bold.

## Discussion

We used GPT-3, a state-of-the art large language model, to show that the speech produced by a large group of first-episode psychosis patients is selectively insensitive to global, relative to local, context. This effect of global-vs.-local insensitivity specifically predicted severity of positive thought disorder (disorganized speech). In contrast, we saw no evidence of a relationship with negative thought disorder (impoverishment) and no relationship with overall symptom severity. These findings have important clinical and theoretical implications.

### Clinical implications: A measure for quantifying positive thought disorder

From a clinical perspective, our index of relative sensitivity to global vs. local context provides an objective, automatic, interpretable measure that can potentially be used to quantify speech disorganization in schizophrenia. The relationship between global-vs-local insensitivity and positive thought disorder was selective. It was also graded: at very local context windows (up to about five words), there was no relationship between lexical probability and positive thought disorder. However, as the window size increased, differences in lexical probability became increasingly large with greater severity of positive thought disorder. In future work, it will be important to extend these findings to patients in the chronic phase of illness and to precisely test the psychometric properties of this measure. With further testing and development, we suggest that it could assist in various clinical purposes, ranging from diagnosis to symptom monitoring. It might also be able to detect subtle language production atypicalities that are less clinically obvious, but that nonetheless impact real-world communicative function. Finally, although the present study focused exclusively on schizophrenia, the methods we have used are very amenable to a transdiagnostic approach, thus laying groundwork for examination of language impairments across diagnostic categories (e.g. bipolar disorder).

### Theoretical implications for a mechanistic understanding of positive thought disorder

Crucially, this measure of global-vs.-local insensitivity goes beyond description. It directly links the clinical phenomenology of thought disorder to neurocognitive constructs that are grounded in psycholinguistic theory.

In healthy adults, numerous psycholinguistic studies of language comprehension have established that healthy adults continually track and use all available contextual information to facilitate lexical processing in real time, as evidenced by both neural (33–38) and behavioral (31,32) measures. In schizophrenia, there is evidence that, although patients’ ability to establish and use local dependencies (world-level priming, clauses, or short sentences) is largely intact (13,16–20), their use of more global sources of information (longer sentences, discourse, or even high-level visual context) during language comprehension is impaired (12–14,16,15). Here, we show, for the first time, that this same global-vs-local insensitivity characterizes natural language *production* in schizophrenia and specifically predicts clinical ratings of positive thought disorder, raising the possibility that atypicalities in language comprehension and production in schizophrenia are driven by shared neurocognitive mechanisms.

These atypicalities are not reducible to the type of generalized cognitive deficits that can impede patients’ performance on challenging neuropsychological tasks (see (71,72)). In healthy adults, the ability to use global context to inform word-by-word language processing is not effortful; rather, it occurs implicitly, without conscious effort. Indeed, in the present study, we found no evidence that patients’ relative insensitivity to global context could be explained by a deficit in overall cognitive function, as indexed by their performance in the neuropsychological tasks we administered. On the other hand, it will be important for future studies to examine the relationship between the atypicalities described here and patients’ performance in tasks that have been linked to selective insensitivities to global context in other cognitive and perceptual domains (e.g., (73–75)) as well as to impairments in the use of context more generally (e.g., (76,77)).

From a computational perspective, selective insensitivity to global linguistic context can be understood within a hierarchical generative framework: a general theory, based on the principles of Bayesian updating, of how brain perceives, interprets and acts upon the world (78,79). Aberrations in hierarchical generative circuits have been proposed as a holistic explanation for multiple neurocognitive atypicalities and symptoms in schizophrenia (80–83), including in language (84,85).

Such a framework naturally explains why the healthy brain is so sensitive to a word’s probability given its prior local and global context (39,86). That is, effective communication requires both the producer and comprehender to employ and continually update hierarchically-organized internal generative models that represent information over successively longer time scales. Individual words (lexical representations) are encoded at relatively short time scales at lower levels of the hierarchy; local semantic/syntactic dependencies are encoded at medium timescales at middle levels; and broader semantic structures (e.g., whole topics and situational contexts) are encoded at the longest timescales at the highest levels of the hierarchy. Thus, within this system, effects of context on lexical probability emerge as a byproduct of optimal communication: as each word is produced/processed in real time, lexical representations serve as a “causal bottleneck”, informed by statistical information encoded at all levels above (31).

In contrast to the human brain, large language models like GPT do not implement probabilistic inference. Rather, they are explicitly trained to predict upcoming words based on vast quantities of human text. Here, we leveraged this property to obtain estimates of lexical probability for each word in patients’ utterances, given local and global context, demonstrating that the disorganized speech produced by first-episode schizophrenia patients fails to benefit from the additional weighting and constraints typically conferred by global information. These findings pinpoint patients’ deficits in linguistic context processing during language production to the highest levels of the generative hierarchy (see (84) for discussion). However, they also raise many additional questions and open up important avenues for future research.

First, what is the precise nature of this high-level atypicality? One possibility is that patients cannot maintain stable representations at the top of their generative hierarchy; that is, at any given time, they may be uncertain about the underlying topic. This type of high-level "belief instability" (87) intuitively explains phenomena that characterize disorganized speech like tangentiality and derailment. Another possibility, however, is that patients fail to flexibly *update* high-level representations (88) (although overly rigid high-level beliefs might also be expected to result in an insensitivity to local context).

GPT is not well-suited for distinguishing these possibilities. Because of its black box nature, we know little about the internal representations it learns. An alternative approach would be to employ language models that are specifically trained to capture latent high-level topic representations from speech outputs (e.g., BERTopic (89)). These topic models could be used to quantify topic diversity within speech samples, allowing us to determine whether uncertainty over topic representations is excessively high in patients’ speech compared to that of healthy controls.

A second set of open questions concerns the specific neurocomputational mechanisms by which atypicalities at the highest levels of representation give rise to selective lexical-level insensitivity to global context during real-time language production and comprehension. Again, most current large language models are ill-suited for addressing this question. Whereas their architectures are feedforward in nature, the human brain is characterized by long-range feedback connections that bridge the highest and lowest levels of the cortical hierarchy. These connections are deeply integrated within the cortical microcircuitry (90), allowing for a continuous interactive exchange of information during word-by-word processing (91).

Understanding how these neural dynamics are affected in schizophrenia would therefore require yet another type of computational model, whose architecture is constrained by what we know about neurobiology. One such model is *predictive coding —* a specific implementation of the more general hierarchical generative framework described above (92–94). Predictive coding is biologically plausible (95–97) and has been able to simulate multiple neural phenomena in human perception, both in healthy adults (92,93,98), as well as in schizophrenia (99).

Indeed, in recent work, we have shown that, in healthy individuals, predictive coding is able to simulate probabilistic effects of context (i.e., effects of lexical probability) on neural activity (100). It can also explain the time course and localization of neural activity produced by incoming words across the left-lateralized fronto-temporal hierarchy during language processing (101). Therefore, an important goal of future studies will be to determine whether the dynamics of predictive coding can explain the specific abnormalities in fronto-temporal neural activity that are commonly observed in thought disorder (for review, see (102)).

### Conclusion

We show that patients in the acute phase of schizophrenia are selectively insensitive to global contextual information during language production. These findings link an objective, theory-driven measure of contextual sensitivity to clinical assessments of positive thought disorder, laying groundwork for an understanding of the computational mechanisms and neural circuitry underlying disorganized language production. Thus, we have begun to realize Bleuler’s original proposal that the basic phenomenology of thought disorder can be linked to the fundamental mechanisms of schizophrenia.

## Acknowledgments

This study was funded by the Sidney Baer Trust. Data acquisition was funded by the Canadian Institutes of Health Research (CIHR) Foundation Grant (Grant no. 375104/2017) to Lena Palaniyappan. Lena Palaniyappan acknowledges research support from the Canada First Research Excellence Fund, awarded to the Healthy Brains, Healthy Lives initiative at McGill University (through New Investigator Supplement to Lena Palaniyappan); Monique H. Bourgeois Chair in Developmental Disorders and Graham Boeckh Foundation (Douglas Research Centre, McGill University) and salary award from the Fonds de recherche du Quebec-Santé (FRQS). We wish to thank Hannah Ke, Sabrina Ford, Cassandra Branco (for data curation and transcription support), Drs. Ross Norman, Peter Liddle (for training support), Drs. Julie Richard, Hooman Ganjavi, Priya Subramanian and Raj Harricharan (for clinical referrals).We are also grateful to undergraduate research assistants Emma Draisin, Brooke Brody, Zoe Gardaret, and Maggie Wargo – and particularly Santiago Noriega, Feng Cheng, and Julia Klein - for their help with data processing and analysis.

## Footnotes

1 The overall study sample had a mean <0.5 defined daily dose equivalents of antipsychotics when speech was assessed (see also (60) for full sample details).

2 We note that the pattern of results was the same (a) when we excluded the five patients with affective psychosis, and (b) when we included the ten patients with missing demographic data.

3 Additional analyses exploring the effects of each cognitive measure separately produced the same results: the interaction between Group and Window Size persisted, whereas there was no main effect, or interaction with Window Size, for any cognitive measure.

