## Supplementary Materials for "GPT-3 reveals selective insensitivity to global *vs.* local linguistic context in speech produced by treatment-naïve patients with positive thought disorder"

Table of Contents

1. Cognitive Tasks
2. Speech Data Collection
3. Thought and Language Index
4. Statistical Analysis
5. Lexical probability in the "NoContext" Condition
6. References

### 1. Cognitive Tasks

In all participants, general cognitive function was assessed using three cognitive tasks. Working memory and processing speed was assessed using a modified version of the Digit-Symbol Substitution Test (DSST; original: (1); modified: (2)), in which participants matched symbols to numbers based on a given key. Both oral and written portions were included, and scores averaged. Semantic memory and executive function were assessed using the Category Fluency task (3) in which participants named as many exemplars of a particular semantic category (e.g. "animals") as possible, and the number of exemplars was tallied. Non-verbal executive function was assessed using Part B of the Trail-Making Test (TMT; (4)) where participants traced a line through consecutive numbers and letters, alternating between numbers and letters (1-A-2-B-3-C-and so on). The time taken to complete this connect-the-dots-style pattern correctly was recorded.

**2. Speech Data Collection**

To obtain the speech data for this study, participants described three pictures from the Thematic Apperception Test (5), for one minute each (see Supplementary Materials for details). Participants were given specific standardized instructions: "I am going to show you some pictures, one at a time. When I put each picture in front of you, I want you to describe the picture to me, as fully as you can. Tell me what you see in the picture, describe what you see in this image, and what you think might be happening." If, for any given picture, the participant stopped speaking before their minute, the experimenter prompted them to go on or give more detail (e.g. "What else do you see?"). Their speech was recorded and transcribed, then used for clinical assessment of thought disorder and extraction of GPT lexical probability measures.

**3. Thought and Language Index**

For each participant, the rater computed a measure of positive thought disorder ("Disorganization" score), which was the sum of ratings for the following TLI items (6): Looseness, Peculiar Word Use, Peculiar Sentence Construction, and Peculiar Logic. They also computed a measure of negative thought disorder ("Impoverishment" score), which was the sum of ratings for the following TLI items: Poverty of Speech and Weakening of Goal.

### 4. Statistical Analysis

We used a series of mixed effects linear regressions to test our primary hypotheses, which were conducted in R (7) using the lmerTest, version 3.1-3 (8) and lme4, version 1.1-33 (9), packages. All continuous predictors were z-scored. The dependent variable (lexical probability) was always log-transformed.

Following the recommendations of Becker (10) and Kalnins (11), we followed up each analysis by repeating the same analysis without nuisance covariates. All significant effects on predictors of interest remained significant after removing covariates, and all non-significant effects of interest remained non-significant.

In all analyses, we modeled the maximal random effects structure justified by the data and our theoretical assumptions. This structure consisted of subject- and, when appropriate, word-level (meaning word token - a particular instance of a word occurring in a discourse segment) intercepts, and random slopes with respect to our predictors of interest when applicable (12). In cases of model non-convergence, if changing the optimizing algorithm did not help, the random effects structure was simplified, beginning with removing correlation parameters for the most complex random effects. All models converged after either changing the optimization algorithm or removing correlation parameters.

**5. Lexical Probability in the "NoContext" Condition**

In our first analysis, in which we tested the broad hypothesis that patients with first-episode psychosis were generally less sensitive to context than healthy controls, we computed baseline measurements of lexical probability, which we term the "NoContext" conditions. To do this, we used GPT-3 (13) to extract lexical probability of the same words, but replacing the the same length of prior context with words that were randomly sampled from an unrelated picture description (based on another picture from the Thematic Apperception Test (5)), which was generated by ChatGPT (see below for prompt and response). The purpose of this measure was to simulate the lexical probability of each given word, free of any relevant context (i.e. no syntactic or discourse structure). That there was no group difference in lexical probability in this condition tells us that any differences in the groups' sensitivity to context could not be attributed to broad differences in the words patients and controls used.

<u>Prompt Given to ChatGPT to Generate Unrelated Picture Description:</u>

*"I am carrying out a study in which I am asking participants to describe black-and-white pictures.*

*The instructions to the participants are as follows:*

*"I want you to describe the picture to me, as fully as you can. Tell me what you see in the picture, describe what you see in this image, and what you think might be happening."*

*I would like you pretend that you are an average participant in this study and you are asked to describe a picture of a boy who is sitting with his elbows on the table and in front of him is a violin and bow. He has his head in his hands and is staring sadly at the violin.*

*Please provide a transcript of a natural utterance that you might produce when asked to describe this picture. You may elaborate on the kinds of things that might be in this picture, but please constrain what you say to what someone might realistically say about a single black and white picture.*

*It should sound like you are actually talking - not writing.*

*The language should sound colloquial without a lot of fancy words - the kind of thing that a regular person might say.*

*The utterance should be 255 words long.”*

Response from ChatGPT:

*“Okay, so there's this black-and-white photo, right? And there's this kid, maybe around 10 or 12 years old, sitting at a wooden table. He's got his elbows on the table and he's leaning forward a bit. He's got short, messy hair, and he's wearing a casual button-up shirt with the sleeves rolled up.*

*Now, right in front of him on the table is a violin and a bow. The violin looks nice and well taken care of. The bow is just lying there next to it, and you can see the strings on it. The black and white in the picture makes everything stand out.*

*The kid has his head in his hands, like he's really sad or something. He's staring at the violin with this downhearted look on his face. It's like he's been crying or is about to cry, you know? It's kinda touching to see.*

*In the background, there's a window with some light coming in. It makes some shadows in the room and adds to the whole sad vibe. You can't really see what's outside, but it might be a cloudy day or something.*

*So, yeah, it looks like the kid is having a tough time with the violin, or maybe something bad happened related to it. The whole thing feels pretty emotional, you know? Like, it really makes you feel for the kid and what he's going through.”*
